# Diversity and distribution of the subtelomeric Y’ elements across *Saccharomyces cerevisiae* strains

**DOI:** 10.1101/2025.11.13.688250

**Authors:** Liébaut Dudragne, Juliana Silva Bernardes, Zhou Xu

## Abstract

The subtelomeric regions of eukaryotic chromosomes harbor repeated elements that contribute to genomic plasticity and adaptation. In *Saccharomyces cerevisiae*, the Y’ elements represent a major class of subtelomeric repeats, yet their diversity and evolutionary dynamics remain incompletely characterized. Here, we analyzed Y’ elements across 54 *S. cerevisiae* strains using high-quality telomere-to-telomere genome assemblies. We detected 893 high-confidence Y’ elements, which we classified into 12 major clusters, revealing a broader structural diversity than previously described, including canonical short (∼5.2 kb) and long (∼6.7 kb) elements, intermediate-size classes (mid1 and mid2), and a novel family containing CA-rich repeats. Sequence analyses showed that open reading frames (ORFs), including those encoding the putative Y’-Help1 helicase, are highly conserved within clusters, suggesting selective maintenance of functional sequences. The distribution of Y’ elements varied widely across strains and chromosome extremities, with some strains lacking Y’ entirely and others, such as the clinical ADI isolate, carrying up to 149 copies. Interstitial telomeric sequences (ITS) were variably associated with Y’ elements and tandem Y’ repeats, potentially facilitating recombination and amplification. Analysis of telomere length data further revealed that the presence of long Yʹ elements, but not Y’ elements from other clusters, at the subtelomere is correlated with shorter telomeres at the same chromosome end. Our results provide the most comprehensive catalog of *S. cerevisiae* Y’ elements to date, uncovering unexpected structural and sequence diversity, and a potentially functional role in telomere length regulation.

## Introduction

Genomic regions located at chromosome extremities are often rich in repeated sequences, including telomeres, the terminal repeats widely studied for their role in genome stability, and subtelomeres, adjacent regions containing diverse families of repeated elements such as transposable elements, satellite sequences, ribosomal DNA and paralogous genes involved in life cycle and adaptative processes (Hocher & Taddei, 2020). Subtelomeres are rapidly evolving regions and display structural and composition diversity even in closely related species or within species, as observed for *Saccharomyces cerevisiae* and *Saccharomyces paradoxus* (Yue et al., 2017). The presence of repeated elements promotes homology-dependent mechanisms, which in turn lead to rearrangements and amplification. In *S. cerevisiae*, the two main types of subtelomeric elements are the Y’ elements and the X elements (Chan et al., 1983; Chan & Tye, 1983). While X elements are found at nearly all extremities, Yʹ elements vary widely in number and organization across subtelomeres and strains, ranging from none to four or more tandem copies often separated by interstitial telomeric sequences (ITS), with some clinical isolates displaying unusually high copy numbers (Chan & Tye, 1983; Heasley & Argueso, 2022; O’Donnell et al., 2023). Y’ elements are usually separated into two types based on their lengths: short Y’ (∼5.2 kb) and long Y’ (∼6.7 kb), differing by an internal deletion (Chan & Tye, 1983; Louis & Haber, 1992), although rare unusually short or long Y’ elements have also been described (Louis & Haber, 1992). Y’ elements have been reported to recombine during mitosis as well as in stress conditions, and to do so preferentially within classes, perhaps explaining how these two classes coexist (Louis & Haber, 1990a).

Y’ elements of previously studied laboratory strains contain a consensus autonomously replicating sequence (ARS), a stretch of 8-20 tandemly repeated 36-mer sequences, as well as one or two open reading frames (ORFs) (Horowitz & Haber, 1984; Louis & Haber, 1992). The ARS sequence seems to be functional, as it is used in *rif1Δ* cells, but does not seem to be used as an active origin of replication in wild-type cells (Theulot et al., 2025). Y’ elements potentially express a DNA helicase called Y’-Help as one of the ORFs contain a motif homologous to eukaryotic helicases and can be transcribed in low amounts (Louis & Haber, 1992; Yamada et al., 1998). Accordingly, most Y’ sequence variations were found to be in the first half of the element, leaving the helicase ORF mostly intact (Louis & Haber, 1992). One study provided further insight by demonstrating that the artificially cloned ORF sequence exhibited functional helicase activity in *Escherichia coli* (Yamada et al., 1998). Despite the presence of this potentially functional ORF, a recent work used an engineered yeast strain containing only three chromosomes to remove all Y’ elements and found no impact on cell growth, telomere length regulation, or telomere silencing (Hu et al., 2024). However, in a specific mutant context where telomere regulator genes are deleted, i.e. *tel1Δ rif1Δ*, Y’ elements exert some effect on telomere length (Craven & Petes, 1999; Sholes et al., 2022). Overall, the physiological functions of the Y’ elements remain elusive.

In telomerase-negative cells, the recombination-based amplification of Y’ elements is the molecular basis of type I post-senescence survivors (Lundblad & Blackburn, 1993; Teng & Zakian, 1999). Y’ elements can be amplified 100 folds in these survivors and invade X-only subtelomeres. The other major senescence bypass pathway, called type II, is characterized by massive telomere elongation, but Y’ elements are also found to be altered and moderately amplified in the early steps of this process (Churikov et al., 2014; Kockler et al., 2021; Teng & Zakian, 1999). Homologous recombination pathways, including break-induced replication (BIR), are required for Y’ acquisition and amplification (Le et al., 1999; Lundblad & Blackburn, 1993; Lydeard et al., 2007). In addition, circular DNA molecules derived from Y’ elements might also contribute to this process (Larrivée & Wellinger, 2006). The Y’-Help helicase is expressed in type I survivors, suggesting it could play a role in subtelomeric recombination (Yamada et al., 1998). Finally, Y’ RNAs can be reversed transcribed by Ty1 retrotransposon to promote recombination in the subtelomeres of telomerase-negative cells (Maxwell et al., 2004).

Beyond their structural diversity and recombination properties, Y’ and X subtelomeric elements also influence chromatin states at chromosome ends. Both contain domains that modulate the telomeric position effect (TPE) (Fourel et al., 1999). Telomere-proximal portions harbor subtelomeric anti-silencing regions (STARs), enriched in Tbf1- and Reb1-binding motifs, and which act as insulators that block the spread of silenced chromatin. In contrast, more centromere-proximal sequences within X and Y’ act as silencing relays, including proto-silencers such as ARS consensus sequences and Abf1-binding sites (Louis, 1995). Such a balance between anti-silencing and silencing activities may explain how Y’ ORFs, such as the Y’-Help helicase, can be expressed despite the generally repressive telomeric environment (Fourel et al., 1999).

Long-read sequencing technologies have recently enabled complete telomere-to-telomere assemblies of *S. cerevisiae* genomes, allowing the accurate resolution of tandemly repeated Yʹ elements in both natural isolates and experimental contexts involving telomere deprotection or erosion (Dudragne et al., 2025; Heasley & Argueso, 2022; Kockler et al., 2021; O’Donnell et al., 2023). Here, we analyze 54 haploid *S. cerevisiae* strains spanning the species’ phylogenetic and ecological diversity to investigate Yʹ element diversity. We describe the most comprehensive repertoire of Yʹ elements to date, uncover a new class containing CA-rich repeats, and characterize the distribution of distinct Yʹ types across chromosome ends and strains.

## Results

### The repertoire of Y’ elements across 54 *S. cerevisiae* strains

As existing Y’ annotations from the *Saccharomyces cerevisiae* Reference Assembly Panel (ScRAP) (O’Donnell et al., 2023) were often fragmented, likely owing to the limited diversity of the single reference sequence used for detection, we developed a bottom-up approach to exhaustively identify full-length Yʹ elements across strains.

Given that telomeric and subtelomeric regions are particularly prone to assembly errors, we focused on the haploid genomes of the ScRAP database to avoid haplotype phasing artifacts inherent to diploid assemblies. Oxford Nanopore Technologies (ONT) sequencing reads were remapped to each assembly for manual inspection and correction of chromosome ends, leading to the manual correction of 35 out of 1696 ends by clipping misassembled terminal sequences. Three strains were discarded due to poor assembly confidence, yielding 53 high-quality haploid genomes for our analysis, to which we added a long-read assembly of the W303 laboratory strain (Garrido et al., 2025). Despite being sequenced as haploids, many of these genomes originate from diploid parental strains, meaning that our dataset captured the genetic diversity present in both haploid and diploid strains. The 54 strains cover a wide range of the diversity of the ScRAP panel, representing 23 out of the 29 clades (Fig. 1A).

**Figure 1.**
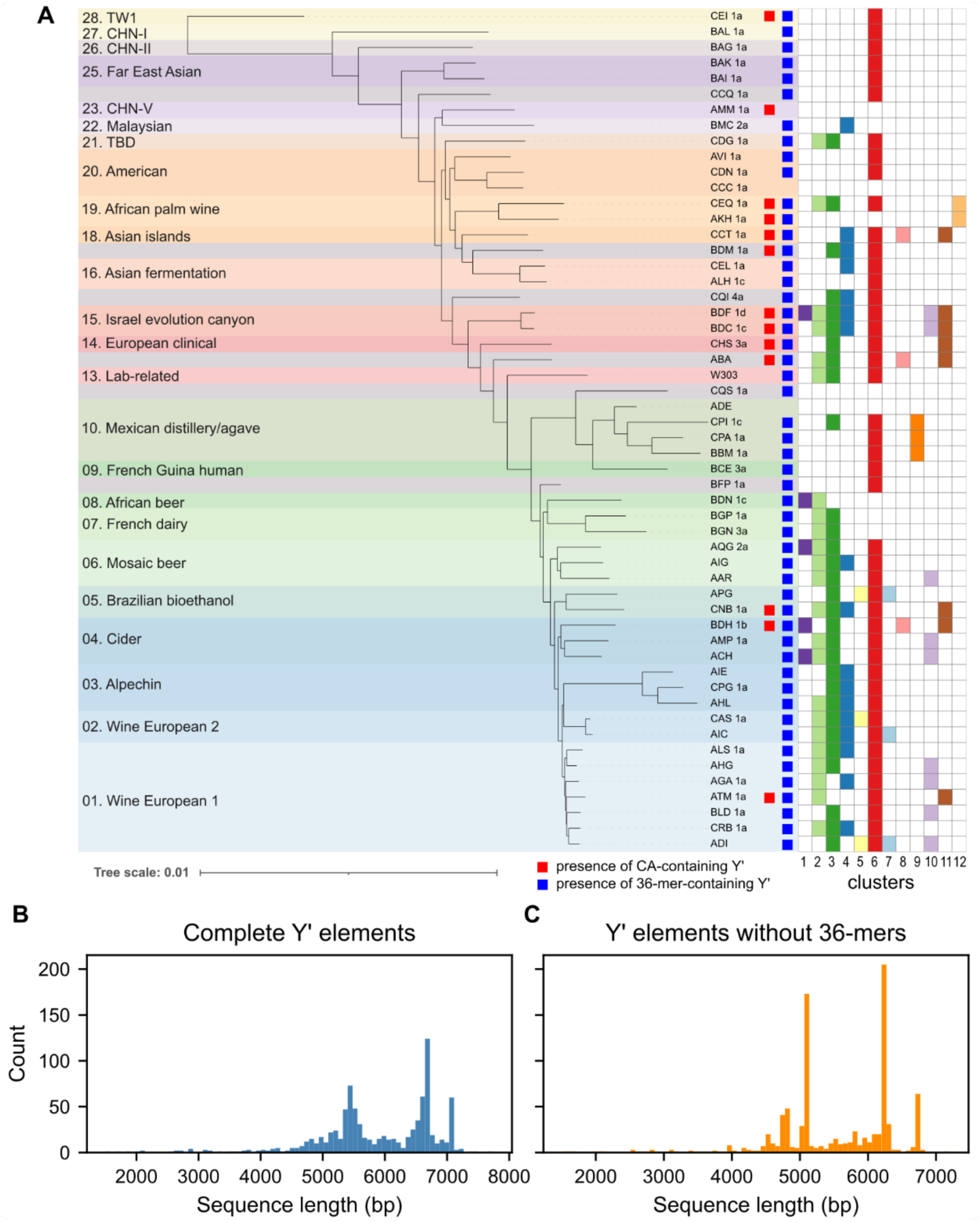
Discrete Y’ element classes with 36-mer repeat variability. (A) Phylogenetic tree of the 54 *S. cerevisiae* strains selected for this study. The tree is a subset of the ScRAP tree extracted from O’Donnell et al., 2023. Colors indicate clades as defined in that same study. Y’ clusters as defined in the next section appear colored when they are present in the strain. (B) Distribution of the lengths of all detected Y’ elements. (C) Same as (B), but after removing the sequence of the 36-mer repeats, revealing sharper peaks as well as new ones.

We sought to build a more complete Y’ reference database for Y’ detection. To establish an initial set of high-confidence, full-length Yʹ sequences, we identified all tandem repeats located in subtelomeric regions and classified as complete Yʹ elements those repeats that matched at least part of the known reference Yʹ element and were directly flanked by telomeric sequences. This led to the detection of 101 high-confidence Y’ sequences, to which we added the Y’ elements of the S288C reference genome. We then mapped these sequences to the 54 genomes with BLAT (Kent, 2002) for exhaustive Y’ detection. This led to the detection of 919 Y’ elements of length >1000 bp, among which 884 elements had a length >3500 bp for a total length of 5,263,491 bp and a mean Y’ length of 5954 bp, which represents an improvement over the 800 Y’ elements for 4,499,540 bp (mean Y’ length of 5624 bp) reported in the ScRAP annotations for the same set of strains. Our method thus successfully detected a greater number of Y’ elements as well as more contiguous and complete copies. 26 elements were removed from further analyses as their coordinates adjoining the extremities of contigs suggested that the assembly was incomplete and the Y’ elements were truncated. Therefore, 893 elements were kept for all downstream analyses (Supp. Data S1 and S2).

### Y’ elements fall into discrete length classes and repetitions of a 36-bp sequence introduce length variability

The lengths of all detected Y’ elements followed a mostly bimodal distribution, with one mode peaking at ∼5,450 bp and another one at ∼6,650 bp, accounting for the two well-known length classes “short” and “long” (Fig. 1B), usually described as being 5.2 and 6.7 kb respectively. In addition, 60 Y’ elements were measured at ∼7,100 bp which, upon further inquiry, were all found to originate from the ADI strain, collected in a clinical environment. The two main modes exhibited substantial spread, suggesting length heterogeneity within the typical length classes. Since Y’ elements have been described as containing a stretch of 36-mer repeats of variable lengths, we asked how much these repeats contributed to the length heterogeneity of Y’ elements. The median length of the stretch of 36-mers was around 360 bp (10 repeats) but ranged from 0 bp (no repeat) up to 1.3 kb (36 repeats) (Supp. Fig. S1A). Analysis of the repeat motif confirmed the consensus sequence found by Horowitz & Haber, 1984 (Supp. Fig. S1B). Interestingly, the variability observed among 36-mer sequences was comparable both within individual Y’ elements and between different Y’ elements (Supp. Fig. S1C), suggesting that sequence degeneration originated in the ancestral Y’ element or that the 36-mer repeats constantly undergo diversification at a high rate.

Removing the 36-mer stretches from the Y’ element sequences reduced the spread around the two main length modes and revealed a new smaller peak below 5 kb (Fig. 1C). Furthermore, a substantial number of elements could still be detected between the “short” and “long” peaks. Together, these findings called for a finer description of Y’ element classes.

### Identification of canonical, intermediate-size, and novel CA-rich clusters of Yʹ elements

To describe the diversity of Y’ elements and identify conserved domains, we calculated a pairwise similarity matrix between all Y’ elements and performed hierarchical clustering, excluding the variable 36-mer repeats which might artificially introduce differences and hinder the analysis (Fig. 2A). It is important to note that the similarity measure, based on per-base match of sequence alignments, is highly sensitive to large insertions and deletions and less sensitive to single nucleotide polymorphisms (SNPs), thus leading to a clustering analysis that prioritizes the structural differences between Y’ elements.

**Figure 2.**
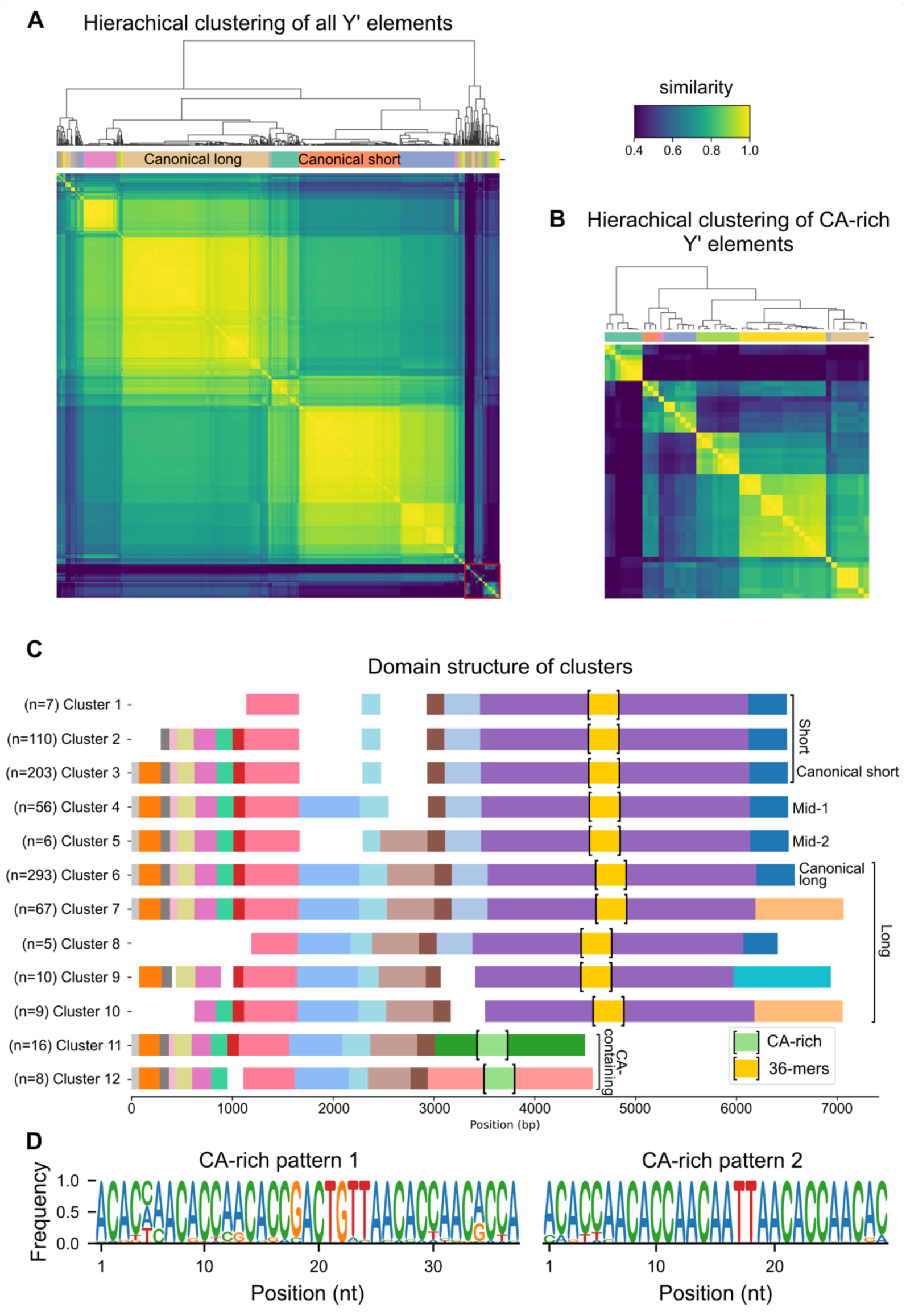
The 12 main clusters of Y’ elements found across *S. cerevisiae* strains. (A) Clustermap of pairwise sequence similarities between Yʹ elements, excluding the 36-mer repeats. The hierarchical clustering tree is shown above. Red square highlights highly divergent sequences. Color scale truncated below the 5^th^ percentile. (B) Clustermap of pairwise sequence similarities between the 49 CA-repeat-containing Y’ elements, computed after removing the CA-rich repeats. The hierarchical clustering tree is shown above. (C) Schematic representation of the 12 main Y’ clusters, using colors to show conserved homology domains. The 36-mer repeats and the CA-rich repeats are depicted between brackets with a fixed length. (D) Logo representation of the two types of patterns found in CA-rich sequences. See Supp. Fig. S2B for per-strain variations of pattern 1.

After optimizing the silhouette score, we identified 46 clusters representing distinct groups of Yʹ elements (Supp. Data S2): the two largest clusters, comprising 293 and 203 elements, corresponded to the canonical long Y’ elements and short Y’ elements, respectively. Together, the five largest clusters accounted for 82% (729/893) of all Y’ elements. Many clusters were very small, containing less than five elements (n = 27) or just one element.

Strikingly, 71 sequences diverged drastically from all others (Fig. 2A, red square at the bottom right of the heatmap). Manual inspection of these elements revealed the presence, in 49 of them, of a stretch of a non-telomeric CA-rich region in the middle of the elements. As for the 36-bp repeats, this CA-rich region was highly variable in length and could artificially introduce noise in our clustering (Supp. Fig. S2A). We therefore removed this region from the Y’ elements before running a similar but separate clustering on these 49 sequences alone, resulting in eight clusters (Fig. 2B and Supp. Data S2).

To focus on the most represented classes of Y’ elements, we dismissed clusters containing fewer than five elements, clusters restricted to a single strain, or clusters with high intracluster heterogeneity. This filter kept 12 clusters containing the vast majority (88%, 790/893) of all Y’ elements for further study. Only 49% (24/49) of CA-containing Y’ elements were kept, indicating that these sequences formed smaller and more heterogeneous clusters.

Using within-cluster multiple sequence alignment, we then retrieved a consensus sequence for each cluster, which we used for further alignment and domain comparison (Fig. 2C). Cluster 6 corresponded to the canonical long Y’ element with a consensus sequence of 6,256 bp, and cluster 3 represented the canonical short Y’ element with a consensus sequence of 5,088 bp, excluding the 36-mer repeats for both. While the short Y’ element differed from the long one by two large deletions of 465 bp and 600 bp, the analysis revealed two clusters (clusters 4 and 5) of Y’ elements containing only one or the other deletion, thus delineating two distinct families of intermediate Y’s that we named Y’-mid1 and Y’-mid2, respectively. Clusters 11 and 12 lacked the second half present in other clusters. Instead, each presented a unique 1.6-1.8 kb domain, within which were nested the CA-rich repeats. Most other clusters differed from the canonical Y’ elements either by having a truncated start or a different end (Fig. 2C), with cluster 7 being almost exclusively composed of Y’ from the ADI strain (64/67 elements).

### Diversity of CA-rich sequences

We examined the CA-rich regions identified within the 49 Y’ elements previously flagged as divergent from canonical sequences. These regions displayed a marked compositional bias, with an average base composition of 45% adenines, 42% cytosines, 7.5% thymines, and 5.5% guanines. This near-equal proportion of adenine and cytosine clearly distinguished them from telomeric CA-rich tracts, which are more enriched in cytosines (62.5% C, 37.5% A).

Inspection of the sequences suggested two recurrent types of repeated motifs, which we characterized by generating consensus sequence logos (Fig. 2D and Supp Fig. S2B). Pattern 1 spans 37 bp, closely matching the length of the canonical 36-mer repeats observed in other Y’ classes. However, no detectable sequence similarity was found between the CA-rich motifs and the 36-mer repeats, suggesting independent origins. Pattern 2, detected in all but 3 strains (CEI, AKH, CEQ), is more variable in length but contains a highly conserved 29-bp core. Its second half bears similarity to the corresponding region of pattern 1, suggesting that pattern 2 may have evolved from pattern 1, or vice versa. Out of all 33,413 bases comprising the extracted CA-rich regions, 19,656 bases (59%) were assigned to pattern 1, and 8,432 bases (25%) to pattern 2.

These observations indicate that CA-rich sequences represent a distinct and heterogeneous component of Y’ elements, with recurrent but diverging motifs that may contribute to structural diversification within this subtelomeric repeat family.

### Distribution of Y’ elements across strains and chromosome extremities

As previously observed (Louis et al., 1994; Louis & Haber, 1990b; O’Donnell et al., 2023), the number of Y’ elements is variable across strains with an average ± standard deviation of 14 ± 9 per strain (Fig. 3A), excluding the ADI outlier displaying 149 Y’ elements. In our dataset, strains ADE and CCC were devoid of Y’ elements and strains BAK, BAI and CCQ contained only one Y’ element, found at chromosome extremity V.L. At the other end of the spectrum, haploid strain ADI presented 149 Y’ elements, spread throughout 25 chromosome extremities. Across all strains, Y’ elements were found at all chromosome extremities (Fig. 3B). Two extremities stood out, with V.L and XIV.R being relatively enriched in Y’ elements. Most chromosome extremities were associated with zero or one Y’ element (1106/1728 and 513/1728, respectively), while tandem repeats of two or more Y’ elements were a relatively rare occurrence (Supp. Fig. S3A).

**Figure 3.**
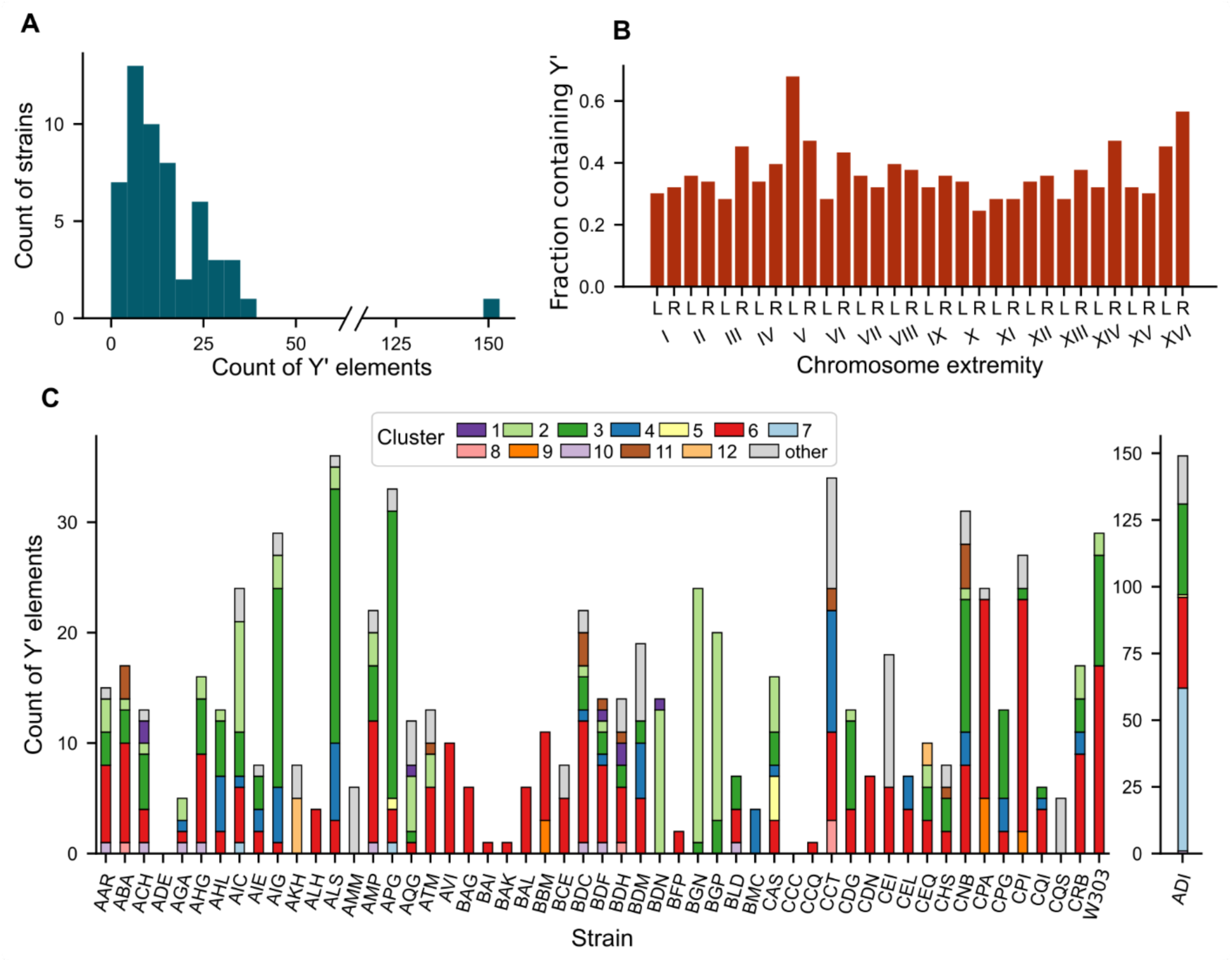
Distribution of Y’ elements across strains and chromosome extremities. (A) Distribution of the number of Y’ elements per strain. (B) Fraction of strains exhibiting at least one Y’ element at corresponding extremities. (C) Distribution of the clusters of Y’ elements across all strains. Strain ADI is set apart with a different scale.

Previous reports suggested that tandem Yʹ elements at a given chromosome end are homogeneously composed of the same length class (Louis & Haber, 1990b; Yamada et al., 1998). However, our analysis showed that this was not always the case: out of the 109 extremities containing tandem Y’ elements, 34 contained a combination of short and long Y’ elements (clusters 1-3 and clusters 6-10, respectively), out of which 25 contained a combination of canonical short and canonical long Y’ elements. Nonetheless, analysis of the conserved regions in Yʹ clusters, excluding the CA-repeat-containing ones, revealed that sequence diversity is lower within strains than between strains, consistent with partial homogenization of Yʹ elements within individual genomes (Supp. Fig. S2B).

We then explored the distribution of Y’ elements per strain according to our clustering analysis (Fig. 3C). Clusters 6 and 3, representing the canonical long and short Y’, respectively, were found in the vast majority of strains (45/54 and 30/54, respectively). Notably, in 9 strains (ALH, AVI, BAG, BAI, BAK, BAL, BFP, and CCQ), the only Yʹ cluster detected corresponded to cluster 6. These strains disproportionately belonged to early divergent lineages (Fig. 1A), and their total number of Yʹ elements was relatively low. Overall, these results depict a scenario in which the canonical long Yʹ of cluster 6 represents the ancestral Yʹ element, from which all other clusters subsequently derived.

Cluster 2, in which the Y’ elements differ from the short Y’ by an additional deletion of ∼300 bp on its left side, is highly represented (>90%) in strains BDN, BGN, and BGP, belonging to two closely related clades: clade 7 (“French dairy”) and clade 8 (“African beer”) (Fig. 1A). Additionally, the vast majority of strains possessing such an element were contained in clades 1-8, suggesting that the emergence of cluster 2 can be traced back to the common ancestor of these clades.

Although CA-containing Y’ elements seemed to be especially present in clades 14 (“European clinical”) and 15 (“Israel evolution canyon”), they were found across the whole phylogenetic tree (Fig. 1A). This suggests that these elements either appeared early in the species and were often lost, or have spread among strains through out-crossing and recombination.

### Open reading frames are highly conserved within clusters and across strains

The presence of ORFs in Y’ elements has been known for a long time, but some discrepancies regarding their length and copy number have been reported between strains and studies (Louis & Haber, 1992; Yamada et al., 1998). Because long-read sequencing is prone to errors, especially in homopolymers, frameshifts in ORFs can sometimes occur in the assembled sequences. Indeed, ORFs were observed to frequently shift reading frames in homopolymers in our Y’ element sequences (Supp. Fig. S4A), although we could not strictly exclude that these frame shifts represented genuine biological events. We quantified the probability of ORF presence at the cluster level by aggregating, across all Yʹ elements, the positions classified as belonging to an ORF, regardless of the reading frame (Fig. 4 and Supp. Fig. S4B). The results clearly indicated a well conserved single long ORF region in the canonical long Y’ cluster (cluster 6) as well as in the mid-1 Y’ cluster (cluster 4), and two smaller well conserved ORF regions in short Y’ clusters (clusters 1, 2 and 3). The first half of the CA-repeat-containing Y’ elements (clusters 11 and 12) also appeared to contain an ORF, in accordance with their high similarity with Y’ elements of other clusters in this region, but in cluster 12, the ORF was split in two parts, whereas in cluster 11, the ORF appeared less conserved, suggestive of a potential evolutionary drift.

**Figure 4.**
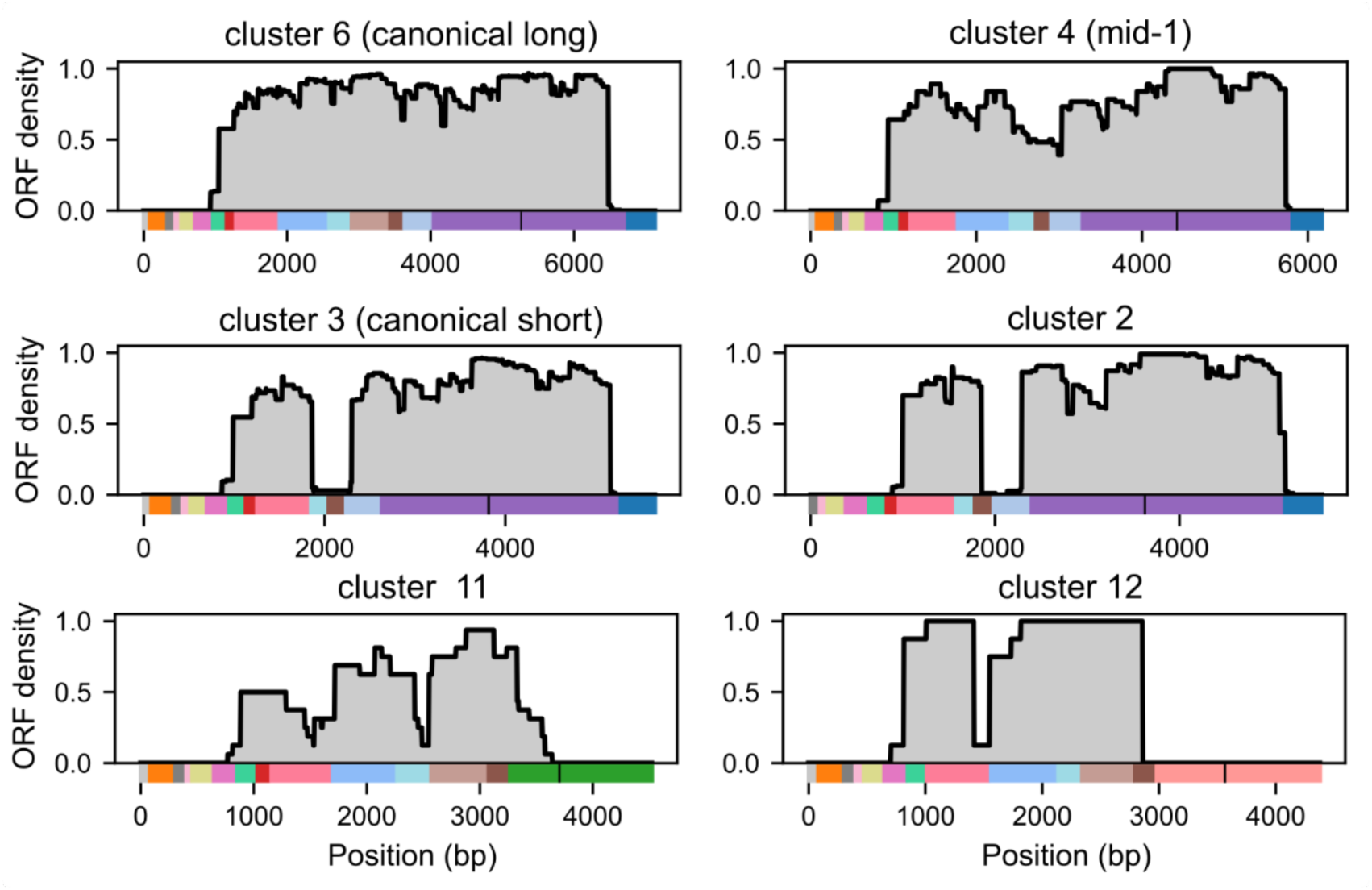
ORF probability within Y’ clusters. Likelihood of a given position in the Y’ element consensus sequence of a cluster belonging to an ORF, computed for 6 clusters. Color bars represent cluster domain structures as depicted in Fig. 2C. Vertical black bars in domain structures indicate the position of the 36-mer or CA-rich repeats. See Supp. Fig. S4B for the other clusters.

### Variability of internal and terminal telomeric sequences associated with Y’ elements

Internal telomeric sequences (ITS) can be found centromere-proximal to Y’ elements and for tandem Y’ repeats, between Y’ elements. Through their homology to the terminal telomere sequence, they contribute to subtelomere rearrangements and Y’ amplification, notably in telomerase-negative cells. We found that only half of the Y’ elements were preceded by an ITS on their centromere-proximal side. Similarly, only 145/234 (62%) tandem repeat junctions presented an ITS in between Y’ elements.

The ITSs displayed a wide length distribution ranging from 20 bp to ∼500 bp and even 628 bp and 957 bp in two instances (Fig. 5A). Overall, the size distributions of ITS between tandem Y’ elements and next to isolated Y’ elements followed a roughly similar shape, with the exception of more very short ITS detected in the proximity (distance < 100 bp) of isolated Y’ elements (Fig. 5A).

**Figure 5.**
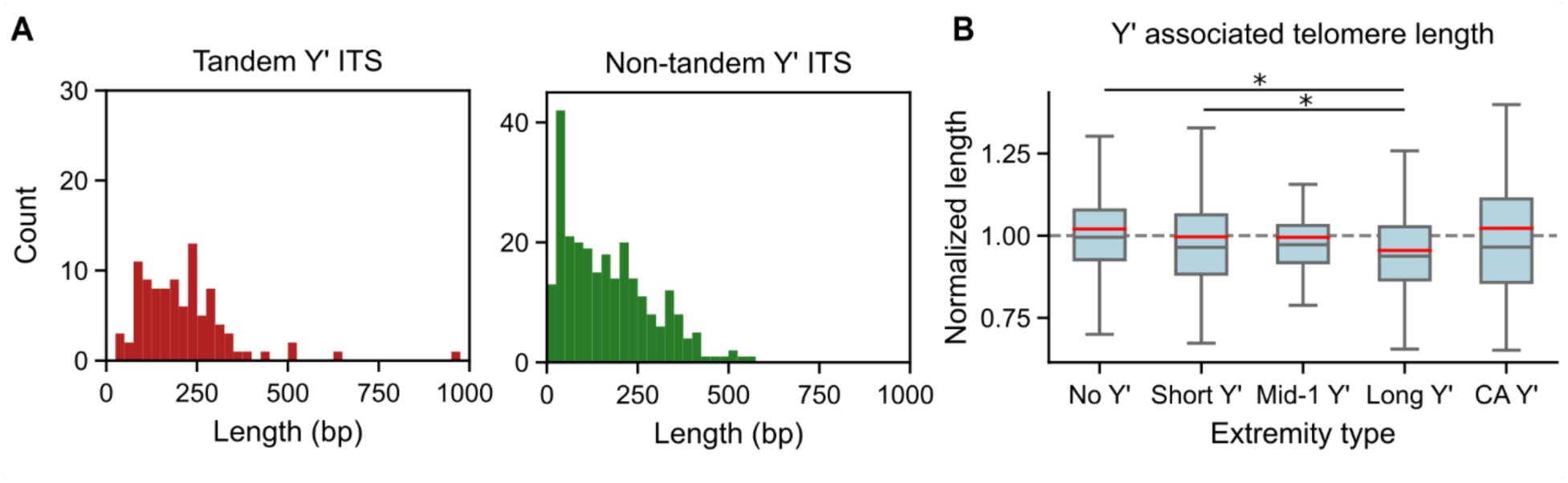
Distributions of ITS lengths and telomere lengths associated with Y’ elements. (A) Distribution of ITS lengths for ITS between tandem Y’ elements (left panel) and not between tandem Y’ elements (right panel), excluding ADI. (B) Boxplot of mean extremity-specific telomere lengths, normalized by the average telomere length in the corresponding strain, associated with different types of Y’ elements. Points outside of the boxplot whiskers are not shown. Red lines indicate mean values. Asterisks show statistical differences with p-values < 0.05 based on two-tailed Student’s t-test.

A recent study reported a subtle but significant influence of Yʹ elements on terminal telomere length: the telomeres immediately adjacent to Yʹ elements are shorter than the ones next to subtelomeres devoid of Y’ element (Garrido et al., 2025). As this analysis was conducted without distinguishing between Y’ types, we reexamined the telomere length data from that study in light of our Y’ classification. We found that telomeres adjacent to canonical long Y’ elements (cluster 6) showed the largest reduction in telomere length: telomeres at these extremities were 6.4% shorter than extremities devoid of Y’ (Fig. 5B), which was significantly more than the impact of short Y’ elements. Telomeres associated with canonical short Y’ element and other short Yʹ elements (clusters 2 and 3) or mid-1 Yʹ elements (cluster 5) were also shorter, but the differences were not individually significant (p = 0.062 and p = 0.304, respectively; Student’s t-test). Combining these classes yielded a statistically significant telomere length reduction (p = 0.042). CA-rich Yʹ elements did not appear to clearly affect telomere length in cis. These results suggest that the region specifically present in canonical long Y’ elements might exert some negative effect on cis telomere regulation.

As Yʹ elements have been reported to bind the silencing regulators Tbf1 and Reb1, we examined whether differences in their binding motifs could account for the distinct effects of Yʹ types on telomere length regulation. However, no differences were observed in the number or position of these binding sites between long and short Yʹ elements, ruling out a role for these regulators in this phenomenon.

## Discussion

In this study, we systematically analyzed hundreds of subtelomeric Y’ elements from strains representing the diversity of *S. cerevisiae* to characterize the structural variations within this family of elements. While 56% of the analyzed Y’ elements fell into what has until now been described as the Y’-short and Y’-long classes, the rest encompassed a large spectrum of structural diversity. The newly identified Y’ classes differed from the canonical ones mostly owing to deletion of domains at the start or the end of the element, sometimes replaced by other sequences. In addition, two distinct repeated motifs (the 36-mer or the CA-rich ones) are found in the second half of the Y’ elements and their variable lengths, up to ∼1.2 or ∼2.1 kb, respectively, add another layer of structural diversity.

The distribution of the clusters across all studied strains suggests that the canonical long Y’ element might have been the ancestral one, from which all others could have derived. The widespread presence of the canonical short Y’ element also suggests that it derived early from the canonical long Y’ and then amplified. Interestingly, we found clusters (mid1 and mid2) with structures that display one or the other of the two deleted parts that differentiate the canonical short Y’ from the canonical long Y’. While any of these two structures could be likely a candidate for an intermediate structure between canonical long Y’ and the canonical short Y’, they are much less abundant and we cannot exclude that they derived independently from the canonical long Y’.

Our analyses revealed a new family of CA-repeat-containing Y’ elements among the most divergent Y’ elements. They are characterized by a CA-rich degenerate repetitive sequence distinct from the 36-mer repeats found in all other Y’ elements. These sequences are highly variable in length and composition, and yet display recurrent motifs (patterns 1 and 2) that are conserved across multiple strains. Intriguingly, in the two main clusters where they are found, the CA-rich repeats are nested within different, non-homologous domains, which are also distinct from the domain found in all other clusters of Y’ elements, thus begging the question of their molecular origin. The near-equal adenine-cytosine content distinguishes these sequences from canonical telomeric CA-rich tracts and suggests that they are not simply divergent telomeric repeats. The strand orientation of the CA-rich repeats also precludes their use as a seed sequence for new telomere addition, should there be an accidental double-strand break occurring close to them. Although none of the identified motifs match the canonical Rap1 binding consensus (Vignais et al., 1990), their similarity to the consensus still leaves open the possibility that Rap1 could bind them directly, albeit with lower affinity.

The number of Y’ elements per strain is widely variable, ranging from 0 to ∼150 in our dataset, highlighting the dynamic and heterogeneous nature of subtelomeres even within a species. We suggest that further investigations on the potential physiological functions of Y’ elements should take into account this natural variability, and a single laboratory strain, such as the W303 strain included in our study, might not be representative. Additionally, it would be interesting to compare strains with highly amplified Y’ elements (e.g. ADI or the strain YJM311 described by Heasley & Argueso, 2022) and telomerase-negative type I post-senescence survivors, with the underlying idea that extended periods of telomere shortening or transient telomere deprotection might have triggered Y’ element amplification (Dudragne et al., 2025; Lundblad & Blackburn, 1993; Teng & Zakian, 1999).

ORFs had previously been identified several times in various Y’ elements (Louis & Haber, 1992; Yamada et al., 1998). However, it remained unclear whether they are functional or merely a vestige of their evolutionary history. While their functionality is still not addressed, we showed that ORFs are well conserved across Y’ elements, with one ORF being present in “long” clusters, and two ORFs being present in “short” clusters, with the exception of CA-repeat-containing Y’ elements showing reduced conservation. This result favors a model in which ORFs are actively conserved within Y’ elements throughout evolution within the *S. cerevisiae* species, suggesting a potentially functional role for Y’ element encoded helicases.

Whether the Y’ element might display a function in telomere length regulation has also been elusive (Craven & Petes, 1999; Hu et al., 2024; Sholes et al., 2022). We recently showed that the presence of Y’ elements is associated with shorter mean telomere length in cis, after normalization by the strain-specific global mean telomere length (Garrido et al., 2025). Here, we refine this observation by showing that long Y’ elements are the main contributor to this association, whereas short or mid1 Y’ elements have no association to shorter telomeres or only a minor one when considered together. We thus speculate that the sequence present in long Y’ elements but absent in mid1 (and short) Y’ elements might recruit a factor that negatively regulates telomere length in cis. Among known factors binding to Y’ elements, both Tbf1 and Reb1 bind to the terminal part of the Y’ element (Fourel et al., 1999) and are not likely candidates.

Overall, this work characterizes the exhaustive repertoire of the highly variable Y’ elements. It uncovers the full spectrum of structural variants, including whole new classes, their distribution across *S. cerevisiae* strains. Beyond the insights it provides on the evolution and potential functions of Y’ elements, this study highlights the diversity and dynamics of subtelomeres, a property common to most eukaryotes.

## Material and Methods

### Data

All data analyzed in this study besides data for W303 strain were downloaded from the European Nucleotide Archive (ENA), from project number PRJEB59869 (O’Donnell et al., 2023). Both reads and assemblies were downloaded, and reads were mapped back on assemblies with Minimap2 (Li, 2018) before visual inspection of chromosome extremities using Tablet (Milne et al., 2013). All chromosome extremities for which there was suspicion of overassembly or misassembly, were trimmed from assemblies. The W303 strain was sequenced and assembled in a previous work (Garrido et al., 2025), and reads and assembly can be found on the ENA under project number PRJEB89249.

### Detection of X elements, Y’ elements and telomeres

Telomeres were detected on assemblies using Telofinder (O’Donnell et al., 2023) (https://github.com/GillesFischerSorbonne/telofinder). This detection was used for terminal telomere detection as well as ITS detection. X elements were detected in the assemblies using LRSDAY 1.7.2 (Yue & Liti, 2018). For Y’ elements, telomeres were first masked from the genomes, then a high-confidence Y’ element database was built by looking for perfect repeats within sliding windows of size 25 kb with a 5 kb step, over the 100 kb extending from every chromosome ends of every strain. Repeats were detected in these windows by mapping the window against itself using BLAST and flagging alignments of lengths comprised between 4.0 and 9.5 kb with an identity percentage of 90 or greater. Yʹ elements from the S288C reference genome (version R64-2-1) (Cherry et al., 2012) were used as queries in a BLAST search across the assemblies, and repeats identified in the previous step that matched at least partially to one of these reference Yʹ sequences were classified as Yʹ elements. To make sure these repeats were detected completely, only repeats directly flanked by telomeric sequences were kept for the high-confidence Y’ element database. This database was then used to detect Y’ elements across all genomes using BLAT v.37×1, with parameter -maxIntron=1000. Because BLAT tends to overextend alignments over large stretches of non-mapping regions, we trimmed the extremal blocks detected by BLAT if they were smaller than 20 bp and the gap separating them from the next block was greater than 100 bp.

### 36-mer detection and masking

All possible 36-mer sequences derived from the degenerate sequence given by Horowitz & Haber, (1984) were used to calculate a 36-meric alignment score across the Y’ elements of the S288C reference genome. At each position, a window of size 36 was extracted, and a score was derived from the Levenshtein distance between the window and the reference 36-mers, calculated as the proportion of matching bases. 36-mers were then detected as peaks in the scores, using scipy.signal find_peaks function with parameters min_distance = 30 and threshold = 0.85. Any match of length different than 36 was discarded, and 36-mers identified this way from the reference genome were used as a new 36-mer database to repeat the same process on every Y’ element from our 54 strains. Matches of size 36 were used to generate a consensus sequence logo using Python’s Logomaker library, and all nucleotides between the first and last detection peak were designated as part of the 36-mer stretch. These regions were then removed from the Y’ element sequences for the subsequent analyses.

### Similarity matrix and hierarchical clustering

Y’ elements were aligned using MAFFT v7.490 (Katoh & Standley, 2013) with “-auto” option, in order to retrieve a multiple sequence alignment (MSA). Similarity matrices were then computed from the alignments by extracting pairs of sequences from the MSA and computing the percentage of shared nucleotides over the full length covered by the union of these two sequences. Positions where the two sequences shared a gap were ignored. The similarity matrix was subjected to hierarchical clustering using seaborn “clustermap*”* function (method = "average"), and clusters were defined by cutting the resulting dendrogram at the chosen cluster count. The optimal number of clusters was defined as the value maximizing the silhouette score across cluster counts from 2 to 100.

The second clustering (Fig. 2B), which ran on CA-containing Y’ elements after the removal of the CA-rich regions, was used to replace the initial cluster assignation of these elements, without changing the cluster assignation of the other elements.

### ORF detection

ORFs were detected as intervals of at least 300 bases between a start codon (ATG) and one of the following stop codons (TAA, TAG or TGA) over the three possible frames. For each element, a masked array indicating the presence of an ORF in any frame was derived. ORF density plots were computed from the masks of all elements of a given cluster, using as a shared coordinate system the coordinates from the MSA of sequences from this cluster.

### Within- vs between-strain diversity of Y’ elements

Y’ elements from clusters 6, 7, 8, 12, 15, 16, 17, 18, 19 and 21, excluding elements from ADI strain, were aligned using MAFFT v7.490 (Katoh & Standley, 2013) with “-auto” option, and the conserved region corresponding to the purple 36-mer-containing region (color from Fig. 2C) was used for diversity analysis. Pairwise distances between elements were then computed from alignment by extracting pairs of sequences from the MSA and computing the percentage of divergent nucleotides. Positions where the two sequences shared a gap were ignored. All pairs of Y’ elements present in the same strain were kept for within-strain boxplot, and all pairs of Y’ elements containing Y’ elements from two different strains were kept for between-strain boxplot.

## Acknowledgement

We thank Gilles Fischer for his critical reading of the manuscript and his feedback. We thank Clotilde Garrido and Oana Ilioaia for their technical help and for discussion. Research in ZX’s laboratory was supported by Agence Nationale de la Recherche (ANR-24-CE12-7740-01 and ANR-24-CE12-0083-03).

## Author contributions

LD: Methodology, Investigation, Conceptualization, Formal Analysis, Writing – Original Draft, Writing – Review & Editing. JSB: Methodology, Investigation, Conceptualization, Formal Analysis, Writing – Original Draft, Writing – Review & Editing. ZX: Conceptualization, Supervision, Writing – Original Draft, Writing – Review & Editing.

## Conflict of interest

The authors declare that they have no conflict of interest.

## Notes

### Competing Interest Statement

The authors have declared no competing interest.

## References

Chan, C. S. M., & Tye, B.-K. (1983). Organization of DNA sequences and replication origins at yeast telomeres. Cell, 33(2), 563–573. 10.1016/0092-8674(83)90437-3

Chan, C. S. M., Tye, B.-K., & Herskowitz, I. (1983). A family of Saccharomyces cerevisiae repetitive autonomously replicating sequences that have very similar genomic environments. Journal of Molecular Biology, 168(3), 505–523. 10.1016/S0022-2836(83)80299-X

Cherry, J. M., Hong, E. L., Amundsen, C., Balakrishnan, R., Binkley, G., Chan, E. T., Christie, K. R., Costanzo, M. C., Dwight, S. S., Engel, S. R., Fisk, D. G., Hirschman, J. E., Hitz, B. C., Karra, K., Krieger, C. J., Miyasato, S. R., Nash, R. S., Park, J., Skrzypek, M. S., … Wong, E. D. (2012). Saccharomyces Genome Database: The genomics resource of budding yeast. Nucleic Acids Research, 40(D1), D700–D705. 10.1093/nar/gkr1029

Churikov, D., Charifi, F., Simon, M.-N., & Géli, V. (2014). Rad59-Facilitated Acquisition of Yʹ Elements by Short Telomeres Delays the Onset of Senescence. PLoS Genetics, 10(11), e1004736. 10.1371/journal.pgen.1004736

Craven, R. J., & Petes, T. D. (1999). Dependence of the Regulation of Telomere Length on the Type of Subtelomeric Repeat in the Yeast Saccharomyces cerevisiae. Genetics, 152(4), 1531–1541. 10.1093/genetics/152.4.1531

Dudragne, L., Garrido, C., Ilioaia, O., Bernardes, J. S., & Xu, Z. (2025). Transient telomere uncapping triggers telomeric and subtelomeric rearrangements. bioRxiv, 2025.07.18.665472. 10.1101/2025.07.18.665472

Fourel, G., Revardel, E., Koering, C. E., & Gilson, É. (1999). Cohabitation of insulators and silencing elements in yeast subtelomeric regions. The EMBO Journal, 18(9), 2522–2537. 10.1093/emboj/18.9.2522

Garrido, C., Kornobis, E., Agier, N., Ilioaia, O., Fischer, G., & Xu, Z. (2025). Natural diversity of telomere length distributions across 100 yeast strains. bioRxiv, 2025.05.13.653712. 10.1101/2025.05.13.653712

Heasley, L. R., & Argueso, J. L. (2022). Genomic characterization of a wild diploid isolate of *Saccharomyces cerevisiae* reveals an extensive and dynamic landscape of structural variation. Genetics, 220(3), iyab193. 10.1093/genetics/iyab193

Hocher, A., & Taddei, A. (2020). Subtelomeres as Specialized Chromatin Domains. BioEssays, 42(5), 1900205. 10.1002/bies.201900205

Horowitz, H., & Haber, J. E. (1984). Subtelomeric regions of yeast chromosomes contain a 36 base-pair tandemly repeated sequence. Nucleic Acids Research, 12(18), 7105–7121. 10.1093/nar/12.18.7105

Hu, C., Zhu, X.-T., He, M.-H., Shao, Y., Qin, Z., Wu, Z.-J., & Zhou, J.-Q. (2024). Elimination of subtelomeric repeat sequences exerts little effect on telomere essential functions in Saccharomyces cerevisiae. eLife, 12, RP91223. 10.7554/eLife.91223

Katoh, K., & Standley, D. M. (2013). MAFFT Multiple Sequence Alignment Software Version 7: Improvements in Performance and Usability. Molecular Biology and Evolution, 30(4), 772–780. 10.1093/molbev/mst010

Kent, W. J. (2002). BLAT —The BLAST -Like Alignment Tool. Genome Research, 12(4), 656–664. 10.1101/gr.229202

Kockler, Z. W., Comeron, J. M., & Malkova, A. (2021). A unified alternative telomere-lengthening pathway in yeast survivor cells. Molecular Cell, 81(8), 1816–1829.e5. 10.1016/j.molcel.2021.02.004

Larrivée, M., & Wellinger, R. J. (2006). Telomerase- and capping-independent yeast survivors with alternate telomere states. Nature Cell Biology, 8(7), 741–747. 10.1038/ncb1429

Le, S., Moore, J. K., Haber, J. E., & Greider, C. W. (1999). RAD50 and RAD51 Define Two Pathways That Collaborate to Maintain Telomeres in the Absence of Telomerase. Genetics, 152(1), 143–152. 10.1093/genetics/152.1.143

Li, H. (2018). Minimap2: Pairwise alignment for nucleotide sequences. Bioinformatics, 34(18), 3094–3100. 10.1093/bioinformatics/bty191

Louis, E. J. (1995). The chromosome ends of *Saccharomyces cerevisiae*. Yeast, 11(16), 1553–1573. 10.1002/yea.320111604

Louis, E. J., & Haber, J. E. (1990a). Mitotic recombination among subtelomeric Y’ repeats in Saccharomyces cerevisiae. Genetics, 124(3), 547–559. 10.1093/genetics/124.3.547

Louis, E. J., & Haber, J. E. (1990b). The subtelomeric Y’ repeat family in Saccharomyces cerevisiae: An experimental system for repeated sequence evolution. Genetics, 124(3), 533–545. 10.1093/genetics/124.3.533

Louis, E. J., & Haber, J. E. (1992). The structure and evolution of subtelomeric Y’ repeats in Saccharomyces cerevisiae. Genetics, 131(3), 559–574. 10.1093/genetics/131.3.559

Louis, E. J., Naumova, E. S., Lee, A., Naumov, G., & Haber, J. E. (1994). The chromosome end in yeast: Its mosaic nature and influence on recombinational dynamics. Genetics, 136(3), 789–802. 10.1093/genetics/136.3.789

Lundblad, V., & Blackburn, E. H. (1993). An alternative pathway for yeast telomere maintenance rescues est1− senescence. Cell, 73(2), 347–360. 10.1016/0092-8674(93)90234-H

Lydeard, J. R., Jain, S., Yamaguchi, M., & Haber, J. E. (2007). Break-induced replication and telomerase-independent telomere maintenance require Pol32. Nature, 448(7155), 820–823. 10.1038/nature06047

Maxwell, P. H., Coombes, C., Kenny, A. E., Lawler, J. F., Boeke, J. D., & Curcio, M. J. (2004). Ty1 Mobilizes Subtelomeric Yʹ Elements in Telomerase-Negative *Saccharomyces cerevisiae* Survivors. Molecular and Cellular Biology, 24(22), 9887–9898. 10.1128/MCB.24.22.9887-9898.2004

Milne, I., Stephen, G., Bayer, M., Cock, P. J. A., Pritchard, L., Cardle, L., Shaw, P. D., & Marshall, D. (2013). Using Tablet for visual exploration of second-generation sequencing data. Briefings in Bioinformatics, 14(2), 193–202. 10.1093/bib/bbs012

O’Donnell, S., Yue, J.-X., Saada, O. A., Agier, N., Caradec, C., Cokelaer, T., De Chiara, M., Delmas, S., Dutreux, F., Fournier, T., Friedrich, A., Kornobis, E., Li, J., Miao, Z., Tattini, L., Schacherer, J., Liti, G., & Fischer, G. (2023). Telomere-to-telomere assemblies of 142 strains characterize the genome structural landscape in Saccharomyces cerevisiae. Nature Genetics, 55(8), 1390–1399. 10.1038/s41588-023-01459-y

Sholes, S. L., Karimian, K., Gershman, A., Kelly, T. J., Timp, W., & Greider, C. W. (2022). Chromosome-specific telomere lengths and the minimal functional telomere revealed by nanopore sequencing. Genome Research, 32(4), 616–628. 10.1101/gr.275868.121

Teng, S.-C., & Zakian, V. A. (1999). Telomere-Telomere Recombination Is an Efficient Bypass Pathway for Telomere Maintenance in *Saccharomyces cerevisiae*. Molecular and Cellular Biology, 19(12), 8083–8093. 10.1128/MCB.19.12.8083

Theulot, B., Tourancheau, A., Simonin Chavignier, E., Jean, E., Arbona, J.-M., Audit, B., Hyrien, O., Lacroix, L., & Le Tallec, B. (2025). Telomere-to-telomere DNA replication timing profiling using single-molecule sequencing with Nanotiming. Nature Communications, 16(1), 242. 10.1038/s41467-024-55520-3

Vignais, M. L., Huet, J., Buhler, J. M., & Sentenac, A. (1990). Contacts between the factor TUF and RPG sequences. Journal of Biological Chemistry, 265(24), 14669–14674. 10.1016/S0021-9258(18)77354-7

Yamada, M., Hayatsu, N., Matsuura, A., & Ishikawa, F. (1998). Yʹ-Help1, a DNA Helicase Encoded by the Yeast Subtelomeric Yʹ Element, Is Induced in Survivors Defective for Telomerase. Journal of Biological Chemistry, 273(50), 33360–33366. 10.1074/jbc.273.50.33360

Yue, J.-X., Li, J., Aigrain, L., Hallin, J., Persson, K., Oliver, K., Bergström, A., Coupland, P., Warringer, J., Lagomarsino, M. C., Fischer, G., Durbin, R., & Liti, G. (2017). Contrasting evolutionary genome dynamics between domesticated and wild yeasts. Nature Genetics, 49(6), 913–924. 10.1038/ng.3847

Yue, J.-X., & Liti, G. (2018). Long-read sequencing data analysis for yeasts. Nature Protocols, 13(6), 1213–1231. 10.1038/nprot.2018.025

