## Supplementary information for "Diversity and distribution of the subtelomeric Y’ elements across *Saccharomyces cerevisiae* strains"

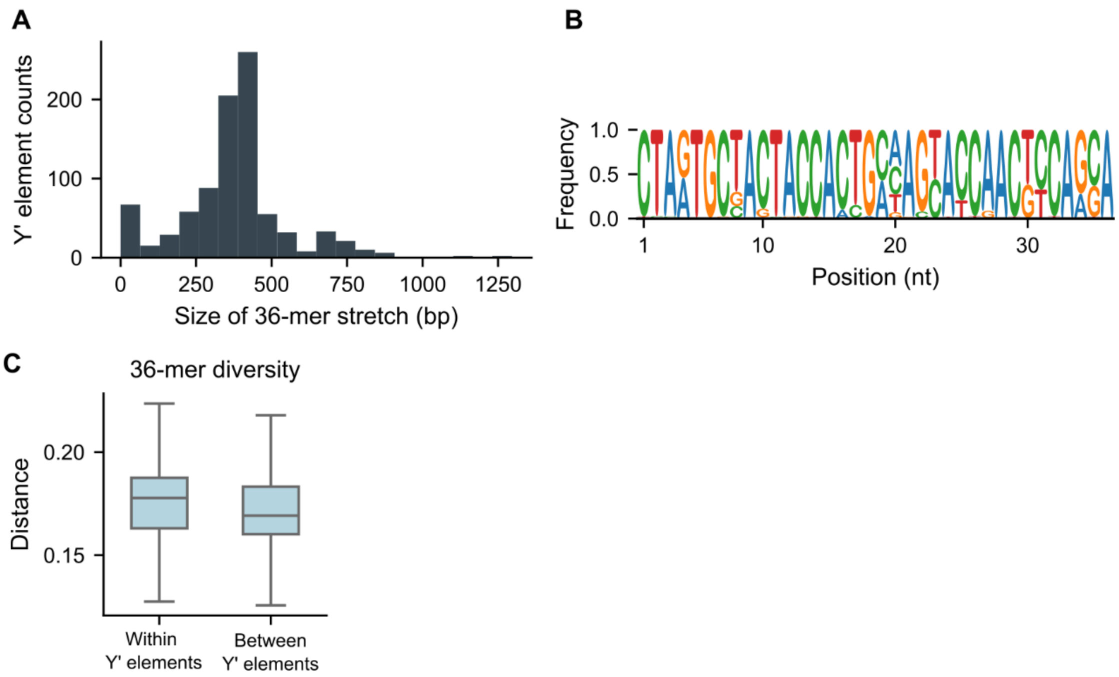


**Supplementary Figure S1. Diversity of 36-mer sequences.**

(A) Distribution of the lengths of the 36-mer repeats.

(B) Logo representation of the consensus sequence of the 36-mer repeats.

(C) Boxplot distribution of pairwise distances of identified 36-mers, within the same stretches and between stretches from different Y’ elements. The points outside of whiskers are not shown.

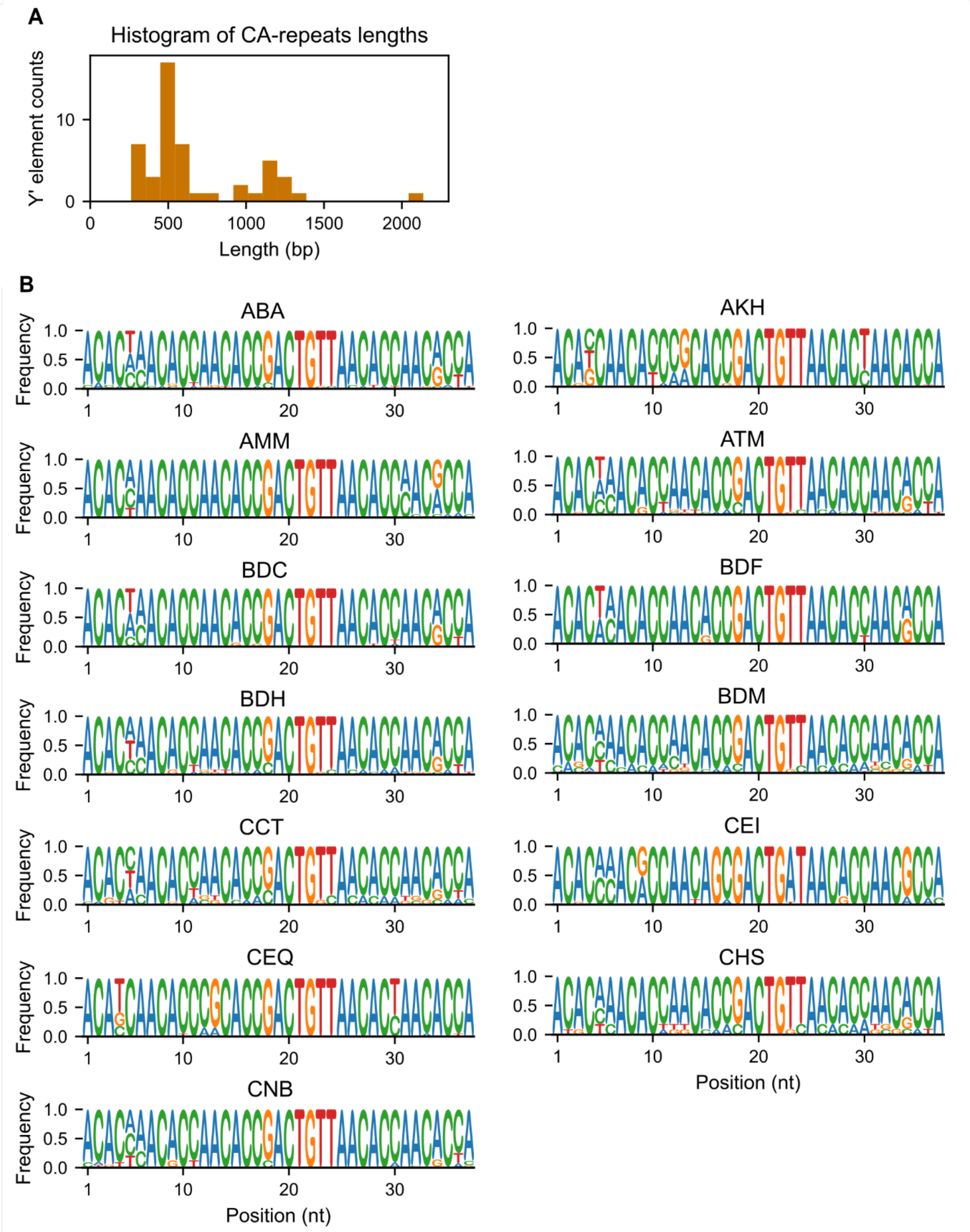


**Supplementary Figure S2. Diversity of CA-rich sequences.**

(A) Histogram of the lengths of the CA-rich regions within the 49 identified CA-containing Y' elements.

(B) Per-strain consensus of CA-rich repeats of pattern 1.

**A**

| Y' count | Extremities |
|---|---|
| 0 | 1042 |
| 1 | 513 |
| 2 | 68 |
| 3 | 15 |
| 4 | 6 |
| 5 | 6 |
| 6 | 6 |
| 7 | 2 |
| 8+ | 6 |

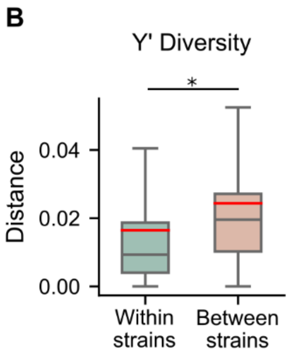


**Supplementary Figure S3. Distribution of Y' elements per extremity and diversity within and between strains.**

(A) Number of tandem Y' elements per chromosome extremity across all strains.

(B) Y' sequence diversity in the conserved purple region of Y' elements from clusters 6, 7, 8, 12, 15, 16, 17, 18, 19 and 21. The points outside of whiskers are not shown. The red lines indicate mean values. *Mann-Whitney U-test, p-value = 0.00.

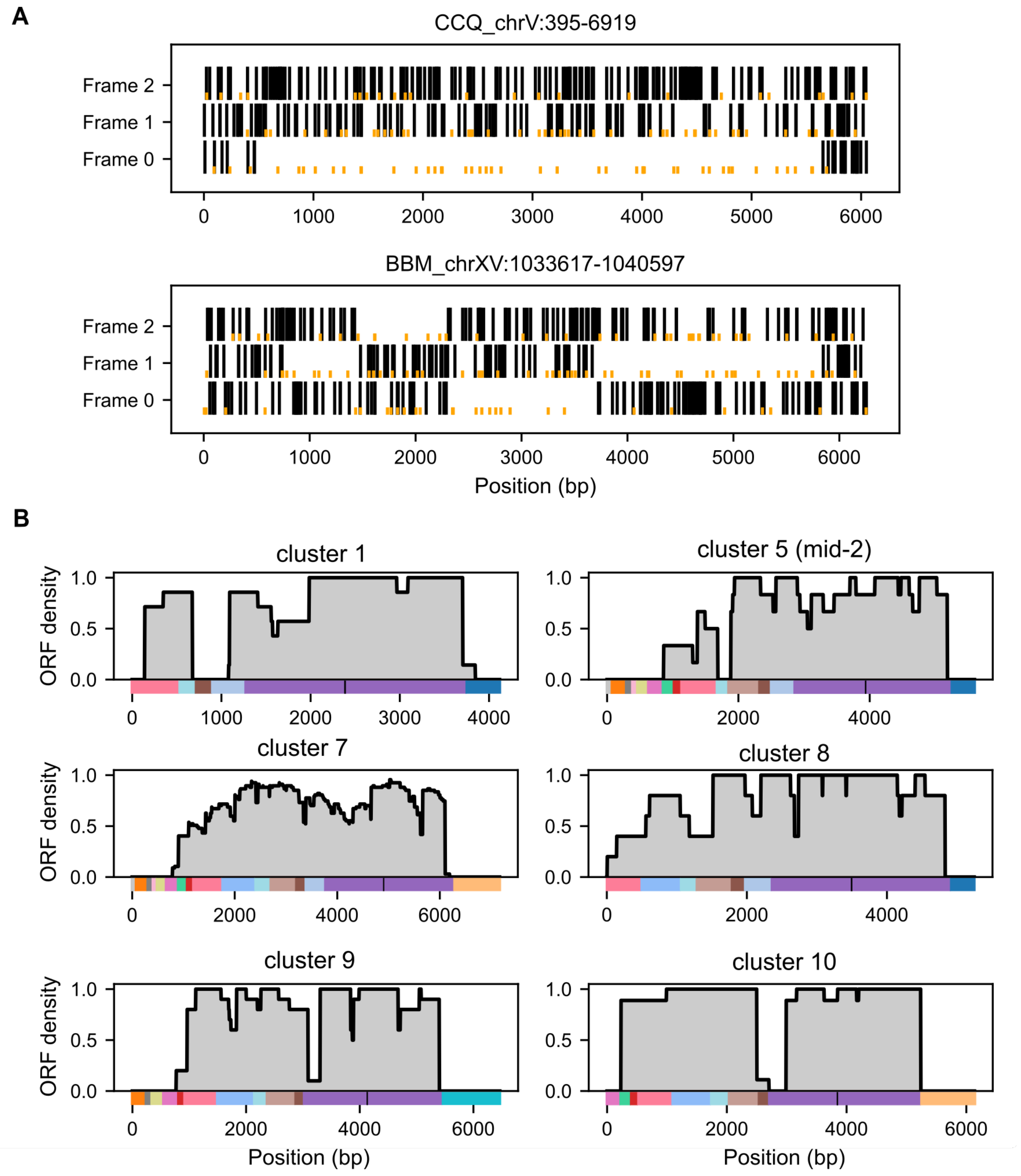


**Supplementary Figure S4. Analysis of the ORFs in Y' elements.**

(A) Start and stop codons across two representative Y' elements allow visualization of potential frame shifts due to sequencing errors. Stop codons are shown as full vertical black lines, start codons are shown as cut vertical orange lines. The id of the Y' elements are shown above each plot. No frameshift event can be seen in CCQ_chrV:395-6919, whereas three frameshifts are observed in BBM_chrXV:1033617-1040597.

(B) Likelihood of a given position in the Y’ element consensus sequence of a cluster belonging to an ORF, computed for the 6 indicated clusters. Color bars represent cluster domain structures as seen in Fig. 2. Vertical black bars in domain structures indicate the position of excised 36-mer or CA streches.

**Supplementary Data S1.** Fasta file containing all 893 Y’ elements detected and analyzed in this work.

The header for each sequence takes the following nomenclature:
StrainName_ChromosomeNumber:start-end.

**Supplementary Data S2.** Table associating each Y’ element, identified by their header in the fasta file in Supp. Data S1, with their cluster number.
